# Not alpha power: prestimulus beta power predicts the magnitude of individual temporal order bias for audiovisual stimuli

**DOI:** 10.1101/2023.06.23.546349

**Authors:** Zeliang Jiang, Lu wang, Xingwei An, Shuang Liu, Erwei Yin, Ye Yan, Dong Ming

## Abstract

Individuals exhibit significant variations in audiovisual temporal order perception. Previous studies have investigated the neural mechanisms underlying these individual differences by analyzing ongoing neural oscillations using stimuli specific to each participant. This study explored whether these effects could extend to different paradigms with the same stimuli across subjects in each paradigm. The two human participants groups performed a temporal order judgment (TOJ) task in two experimental paradigms while recording EEG. One is the beep-flash paradigm, while the other is the stream-bounce paradigm. We focused on the correlation between individual temporal order bias (i.e., point of subjective simultaneity (PSS)) and spontaneous neural oscillations. In addition, we also explored whether the frontal cortex could modulate the correlation through a simple mediation model. We found that the beta band power in the auditory cortex could negatively predict the individual’s PSS in the beep-flash paradigm. Similarly, the same effects were observed in the visual cortex during the stream-bounce paradigm. Furthermore, the frontal cortex could influence the power in the sensory cortex and further shape the individual’s PSS. These results suggested that the individual’s PSS was modulated by auditory or visual cortical excitability depending on the experimental stimuli. The frontal cortex could shape the relation between sensory cortical excitability and the individual’s PSS in a top-down manner. In conclusion, our findings indicated that the prefrontal cortex could effectively regulate an individual’s temporal order bias, providing insights into audiovisual temporal order perception mechanisms and potential interventions for modulating temporal perception.

## 1 Introduction

We reside within a multimodal world where the integration of our senses hinges upon our ability to perceive and discern the precise temporal relationships between diverse modal stimuli. Cross-modal temporal perception is crucial for multisensory integration and high-level cognition ^[1-3]^. However, multisensory temporal perception is seldom veridical and characterized by substantial interindividual variability ^[4, 5]^.

Undoubtedly, the differences in the processing capacity of sensory information contribute to variations in temporal perception, including local neural oscillations ^[6, 7]^ and the interactions among neural populations ^[8, 9]^. For instance, subjects with narrower temporal binding windows (TBWs) exhibited a more detailed neural representation of the sound onset ^[6]^. The increased interactions of temporal and frontal cortical regions, associated with multisensory integration and response selection, are related to higher temporal sensitivity ^[8]^.

Spontaneous neural activities also modulate temporal perception ^[10-13]^. Using a simultaneity judgment (SJ) task, we found that prestimulus alpha power in the visual cortex positively predicts the magnitude of an individual’s Point of Subjective Simultaneity (PSS) ^[12, 13]^. Prestimulus alpha power in the auditory cortex is also positively associated with an individual’s PSS with a temporal order (TOJ) task ^[11]^. Furtherly, when the authors matched two groups of subjects whose PSSs are in opposite audiovisual temporal sequences (i.e., auditory first or visual first), they found the frontal alpha band power positively correlates with the magnitude of PSS ^[10]^. The alpha power in the auditory or visual cortex reflects specific cortical excitability: higher alpha power corresponds to lower cortical excitability and poorer perceptual processing ^[14]^. In contrast, the alpha power in the frontal cortex reflects top-down cognitive control ^[10]^.

After carefully examining the experimental designs of the two papers with the TOJ task ^[10, 11]^, we inferred that the experiment’s stimulus settings might reveal different neural mechanisms in reflecting the variations of an individual’s temporal perception. Grabot et al.(2017) extracted the subjects’ specific PSSs, JND1 and JND2 (just noticeable difference), for EEG experiments. In statistical analysis, they performed Pearson’s correlation between the relative power (Visual first minus auditory first response; Vfirst-Afirst) and PSS in PSS-related stimulus onset asynchrony (SOA) conditions ^[11]^. In contrast, Grabot et al.(2020) derived only the subjects’ specific JND1 and JND2 for EEG experiments. They performed Pearson’s correlation analysis in JND1 and JND2-related SOA conditions ^[10]^. So we could conclude that when participants are exposed to more challenging stimuli, their perceptual responses are modulated by the sensory cortex. Conversely, when encountering relatively simple stimuli, the higher-level prefrontal cortex primarily regulates their perceptual responses.

As PSS is derived from the fitting curves with multiple stimulus onset asynchrony (SOA) conditions, it might be affected by the cortical states across all SOA conditions. Therefore, we aimed to explore whether the prestimulus cortical states across all SOA conditions predict the magnitude of an individual’s PSS. Based on previous experimental designs and results, we guessed that prestimulus sensory cortical excitability might affect an individual’s PSS. The reason is that when the subject faced multiple SOA conditions, the participants might perceive all conditions as relatively challenging as the varying difficulty levels of the stimuli throughout the experiments.

Recent research indicated that sensory cortical excitability is a bottom-up process that can be modulated by top-down signals from the higher-level frontal cortex ^[15-17]^. Interregional oscillatory synchrony might be a candidate mechanism. A study found that frontal-occipital phase synchronization predicts occipital alpha power in visual perceptual decision-making ^[15]^. Furthermore, combining TMS, EEG, and behavioral tasks, recent research has indicated that the frontal eye field can modulate the excitability of the visual cortex, thereby causally influencing visual perception through mechanisms of oscillatory realignment measured by phase resetting ^[17]^. If the prestimulus sensory cortical excitability affects an individual’s PSS, we wanted to explore whether the frontal lobe regulates sensory cortical excitability, thereby modulating individual PSSs.

In this study, we utilized a novel experimental design, encompassing all SOA conditions and ensuring consistency across subjects, to investigate the neural mechanisms underlying individual variability in PSSs. We analyzed two distinct EEG datasets. The first dataset (dataset A), publicly available, involved subjects performing a TOJ task with simple auditory beeps and visual flashes as stimuli ^[18]^. The second dataset (dataset B) we collected involved participants performing the same task using the streaming/bouncing illusion paradigm. Our results indicated that prestimulus sensory cortical excitability negatively predicted the individual’s PSS. Notably, the spontaneous auditory cortical excitability is related to the beep-flash paradigm, while visual cortical excitability is correlated with the streaming/bouncing illusion paradigm. Importantly, we found that frontotemporal or frontal-occipital interregional interactions measured by phase synchronization shaped the prestimulus auditory or visual cortical excitability, further affecting the individual’s PSS. These findings suggested that sensory cortical excitability reflected the individual’s PSS depending on the experimental paradigm and specified the general link between frontal top-down modulation, sensory cortical excitability, and performance in perceptual tasks.

## 2 Material and methods

### 2.1 Participants

In dataset A, forty-three subjects participated in the experiment. Three participants were excluded due to bad behavioral results. Forty participants (30 female, two left-handed, median age: 23, age range:18–32) remained in data analysis. The Ethics Committee of Ghent University approved the experiment.

In dataset B, we recruited thirty subjects to participate in the experiment. One participant was excluded due to poor EEG data. Twenty-nine participants (11 female, all right-handed, median age: 21, age range:17–25) remained for statistical analysis finally. The Tianjin University Institutional Review Board approved all recruitment and experimental procedures.

In two datasets, all the subjects had normal or corrected to normal vision and normal hearing and no history of neurological disorder.

### 2.2 Stimuli

In dataset A, the visual stimulus was a solid white circle with a 1.95° visual angle with a duration of 16 ms. The auditory stimulus was an 1850-Hz tone presented for 16 ms (see Fig. 1a). Please refer to the paper for more detailed information ^[18]^.

**Fig. 1.**
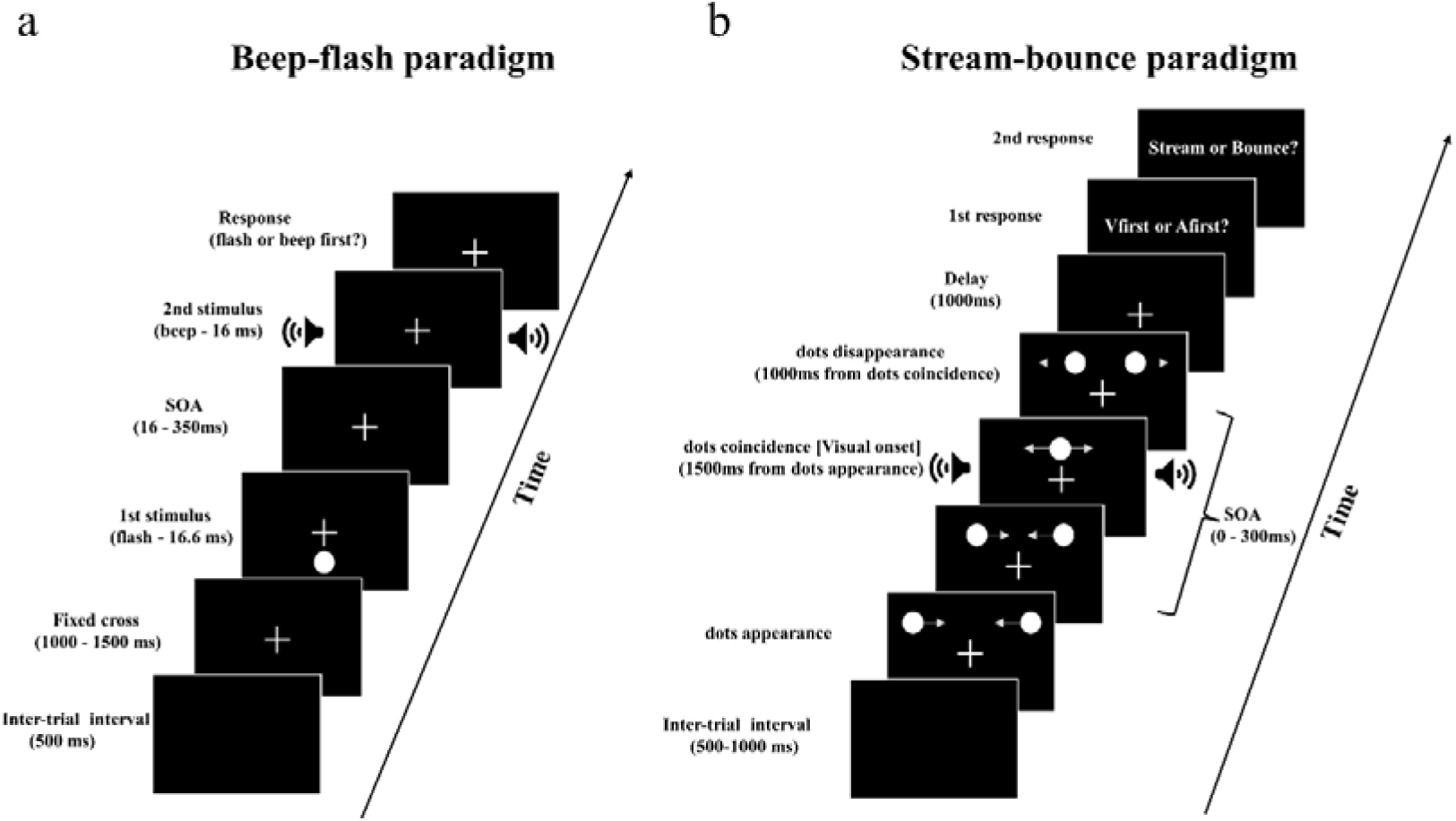
Schematic illustration of the experimental paradigms. Example of a single trial in the beep-flash (a) and stream-bounce (b) paradigm.

In dataset B, the visual stimuli were two white dots (a diameter of 1°) presented on a 23-in monitor (Philips 236V6Q) (refresh rate= 60 Hz, resolution = 1920 ×1080). The auditory stimulus was an 800 Hz pure tone lasting 50 ms with 3 ms rise and fall ramps (see Fig. 1b).

### 2.3 Experimental design and procedure

The experimental designs of both datasets are single-factor within-subject designs. Each participant completed the TOJ task with twelve different Stimulus Onset Asynchronies (SOAs) [± 350, ± 216, ± 133, ± 88, ± 50, ± 16 ms] in dataset A, while the subject with seven different SOAs (± 300, ± 200, ± 100, and 0 ms) in dataset B. Negative SOAs indicated the auditory stimuli were first presented (AV). Positive SOAs implied the visual stimuli appeared first (VA). The SOA with 0 ms means that the visual and auditory stimuli appeared simultaneously. Notably, we defined two dots’ point of coincidence (POC) as the visual onset ^[19]^ in dataset B. Each SOA was repeated 70 times. The 840 trials were divided into 35 blocks with two repetitions for each SOA in dataset A. Each SOA was repeated 40 times. The 280 trials were divided into eight blocks with five repetitions for each SOA in dataset B. Four subjects performed twelve blocks with 420 trials in dataset B.

The task ran on an HP Compaq desktop computer with the E-prime 1.2 software package in dataset A. The experimental process was as follows: First, an inter-trial interval would last 500 ms. After that, a fixation cross would appear on the screen lasting for a random time ranging from 1000 to 1500 ms, followed by a pair of beep-flash stimuli with a random SOA. The fixation cross disappeared after participants had answered the flash or beep first (Vfirst or Afirst) (see Fig. 1a). Participants pressed the ‘z’ key when they had perceived Afirst and the ‘m’ key when they had perceived Vfirst.

The task ran on a Philips desktop computer in dataset B with the Psychophysics Toolbox 3.0.11 and MATLAB R2017a (MathWorks). The experimental procedure was outlined as follows. After an inter-trial interval of 500-1000 ms, two dots appear simultaneously on both sides of the screen with a fixation cross on the center. The two dots started to move horizontally on the screen at a velocity of 0.19° per frame. After 1500 ms, the two dots converge together above the center of the screen. Then the two dots continued moving for 1000 ms. After the dots disappeared, a response delay would appear to prevent the subjects’ motor preparation signal from interfering with the EEG signals we focused on. Finally, two response interfaces appeared. The subjects made a temporal order response: Vfirst or Afirst. Then they should make another response: stream or bounce (see Fig. 1b). Participants pressed the ‘5’ key when they had perceived Afirst and the ‘4’ key when they had perceived Vfirst. They made a stream response with the ‘1’ key and a bounce response with the ‘2’ key. Note that we only focused on temporal order responses in this paper.

### 2.4 EEG recording and preprocessing

The electroencephalogram (EEG) was recorded at 1024 Hz in dataset A using a Biosemi ActiveTwo system (Biosemi, Amsterdam, Netherlands) with 64 Ag–AgCl scalp electrodes positioned according to the standard international 10–20 system. Notably, there existed only the clean data after preprocessing. The EEG of dataset B was continuously recorded using a SynAmps amplifier (Neuroscan) with 62 Ag/AgCl electrodes positioned according to the international 10–20 system. The sampling rate was set at 1000 Hz, and the impedance of each electrode was maintained below 5 KΩ. Online referencing against the left mastoid was used.

We employed a similar data preprocessing approach in dataset B as in dataset A to ensure comparability between the two datasets. Data were first high-pass and low-pass filtered at 0.5 and 30 Hz, separately. Then the data was cut into 2 s epochs (−1500 ms from the first stimulus onset to 500 ms). The data were then re-referenced to the average of all electrodes in dataset A. The EEG channels were firstly re-referenced to two linked electrodes on the left and right mastoid and then to the average of all remaining 60 electrodes in dataset B. We used Independent Component Analysis (ICA) to remove artifacts. The sampling rates were resampled to 200 Hz. Finally, the Laplacian transform was applied to improve topographic localization. All subsequent data processing was conducted in Matlab (MathWorks) utilizing the EEGLAB ^[20]^ and Fieldtrip ^[21]^ Toolbox.

### 2.5 Behavioral analysis

We used the percentage of Vfirst responses across all SOA conditions throughout the experiment to fit a logistic curve by the *glmfit* function in Matlab to determine the temporal order bias [point of subjective simultaneity (PSS, the point with ∼50 % proportion of Vfirst responses among all SOA conditions)] for each participant using the method of constant stimuli ^[22]^. The goodness of fit (R^2^) was 92.59 ± 7.06 % and 93.75 ± 4.69 % on average across participants (mean ± SD) for datasets A and B, respectively. In datasets A and B, the goodness of fit for all participants were above 85%. One participant was excluded from the formal analysis as the goodness of fit is 60.39 % in datasets A.

To investigate the stability of PSS across time, we re-fitted the logistic curve for each participant based on 20 trials at each SOA. The 35 blocks were divided into six groups with five block steps in dataset A (see Fig. 2c and d). For example, the first subgroup is from block 1 to block 9. The second subgroup is from block 5 to block 12. The eight blocks were divided into three groups with two block steps in dataset B (see Fig. 2e and f). For example, the first subgroup is from block 1 to block 4. The second subgroup is from block 3 to block 6. We defined the first subgroup in dataset A as GroupA1 while in dataset B as GroupB1. The GroupAall was defined as a PSS group including all trials of the six subgroups in dataset A, while GroupBall in dataset B. We excluded any participants from each group whose goodness of fit was below 75%. There were 33 remaining participants in dataset A, while there were 24 remaining participants in dataset B. The final goodness of fit in each subgroup is in table one.

**Fig. 2.**
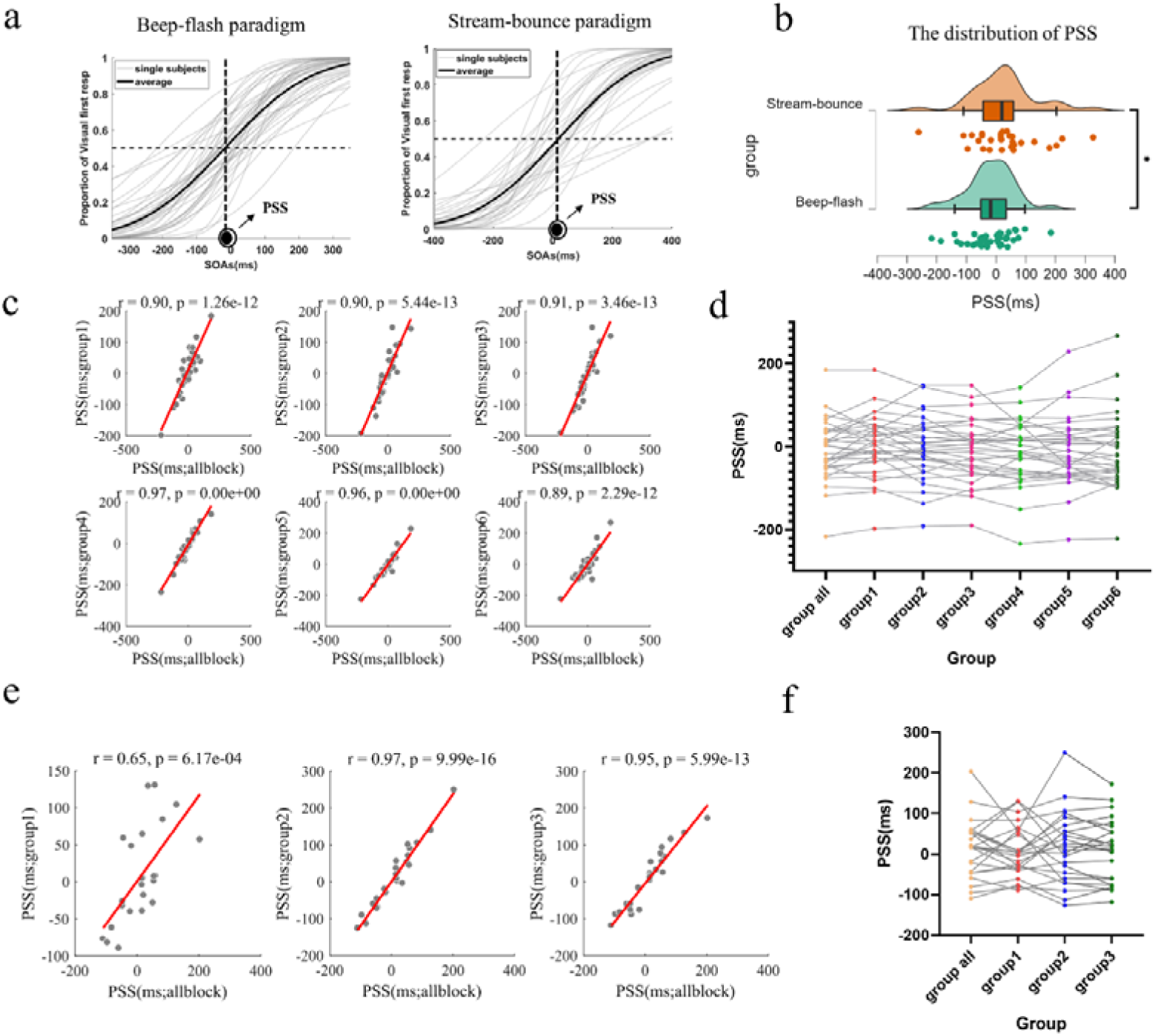
The results of behavior responses in two experimental paradigms. (a) Schematics of the psychometric curves for each participant (in grey) and the group-averaged value (in dark black). Negative values indicate auditory leading, while positive values indicate visual leading. The PSS (Point of Subjective Simultaneity) corresponds to the horizontal axis SOA at which the 50% proportion of visual first responses was observed. (b) The Distribution of PSSs under two experimental paradigms. * p < 0.05. (c) Pearson’s correlations were computed between the six subgroups of PSSs and the PSSs derived from all trials for each participant in the beep-flash paradigm. Note that the sample size is 33 after deleting the subjects whose goodness of fit was below 75% in any of the seven subgroups. (d) The distribution of PSSs in each subgroup in the beep-flash paradigm. (e) Pearson’s correlations were computed between the three groups of PSSs and the PSSs derived from all trials for each participant in the stream-bounce paradigm. Note that the sample size is 24 after deleting the subjects whose goodness of fit was below 75% in any of the four subgroups. (f) The distribution of PSSs in each subgroup in the stream-bounce paradigm.

**Table 1.**
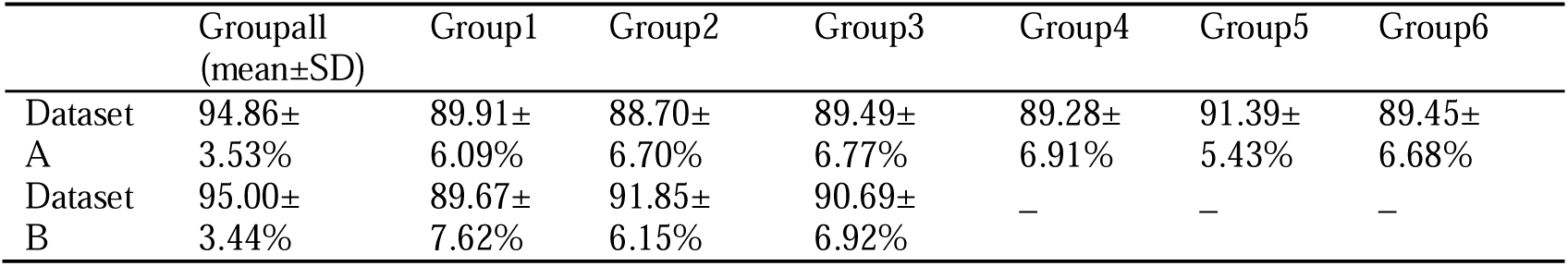
The goodness of fit for the sub-group in datasets A and B.

### 2.6 EEG analysis

#### 2.6.1 Time-frequency decomposition

The single trial’s complex time-frequency and cross spectra were extracted by Morlet wavelets with fixed seven cycles between 1 and 30 Hz with a step size of 1 Hz from −1.5 s to 0.5 s around the first stimulus onset. The amplitude of poststimulus EEG was set to zero to avoid any poststimulus contamination on spontaneous neural activities ^[10]^. The time period of interest is from −500 ms to 0 ms in statistical analysis.

#### 2.6.2 Spontaneous oscillatory power analysis

The raw power was calculated by computing the squared magnitude of the complex time-frequency spectra and log-transformed to control for potential influences of non-normality for statistics ^[23]^.

#### 2.6.3 The dwPLI analysis

Phase synchrony is one of the primary ways to quantify functional connectivity between regions. We used the debiased weighted phase lag index (dWPLI) ^[24]^, used in many related papers ^[15, 25]^, to quantify phase synchronization. The dWPLI is not sensitive to volume conduction and more appropriate for scalp-level connectivity ^[26]^. We calculated the dwPLI metrics with the function “ft_connectivity_wpli.m” in the Fieldtrip toolbox.

### 2.7 Statistical analysis

#### 2.7.1 Behavioral data statistical analysis

First, we used one-sample t-tests to verify whether the PSSs in the two datasets were significantly different from zero. The independent samples t-test was to used to test the difference between PSSs in two datasets. Then, we performed Pearson’s correlation analysis between the PSSs of Groupall and each subgroup in two datasets. Finally, we used one-way repeated measures analysis of variance (ANOVA) to investigate the PSSs difference between each subgroup.

As we focused on power and dwPLI difference between auditory and visual first responses in EEG analysis, we used paired sample t-test to compare the trial numbers between the Vfirst and Afirst responses to exclude the influence of trial numbers on the evaluation of power and dwPLI difference.

#### 2.7.2 EEG statistical analysis

We performed the Pearson’s correlation between the power difference (Afirst minus Vfirst; Afirst-Vfirst) and the individual’s PSS using the ft_statfun_correlationT function in the Fieldtrip toolbox. Then the cluster-based permutation test was used to control multiple comparisons. The calculation process is as follows. First, we calculated Pearson’s correlation for power difference and the individual’s PSS. Data points with p-values less than 0.05 are selected, and clusters are formed based on adjacency between temporal, spectral, and spatial domains. We set the spatial adjacency with at least two adjacent channels. Then, we chose the cluster with the maximum or minimum cluster values to represent the empirical-level statistics. Second, Monte-Carlo randomization 1000 times was performed to shuffle the labels of the participants and completed the Pearson’s correlation again. We could get the 1000 clusters as a surrogate distribution. The empirical-level statistics were considered significant if they were higher or lower than the surrogate distribution’s upper 97.5th percentile or lower 2.5th percentile.

The same process was used to assess the significance of Pearson’s correlation between the dwPLI difference (Afirst minus Vfirst; Afirst-Vfirst) and the individual’s PSS or power difference (Afirst-Vfirst). Note the clusters were only based on temporal and spectral adjacency.

We used the mediation model to reveal the relationship between dwPLI, power, and the individual’s PSS. The PROCESS macro in SPSS was used for mediation analysis. The significance of the path coefficients in the mediation model was assessed with the bootstrap method. The path coefficients’ 95% bootstrap confidence intervals (CIs) were calculated based on 5000 bootstrap samples.

## 3 Results

### 3.1 Behavior Results

We used one-sample t-tests to test whether the mean of the individual’s PSS was about 0 ms in the two datasets. The results indicated that there was no significant difference against 0 ms at the two datasets [dataset A: −18.038 ± 77.784 (mean ± SD), *t* _(38)_ = −1.448, *p* = 0.156; dataset B: 28.590 ± 113.048 (mean ± SD), *t* _(28)_ = 1.362, *p* = 0.184]. The results of the paired-sample t-test showed that the mean of the individual’s PSS in dataset A was significantly smaller than in dataset B [*t*_(66)_ = −2.015, *p* = 0.048, Cohen’s d = −0.494, Fig. 2b]. We calculated the Pearson’s correlation between the PSS obtained from fitting all trials and the PSS obtained from trials in the subgroup to test the stability of PSS across time. The results showed that all correlation coefficients were significant (Fig. 2c and e). Further, repeated-measures one-way ANOVA revealed that there was no significant effect of time on PSSs [dataset A: *F* _(6, 266)_ = 1.299, *p* = 0.259, η*p^2^* = 0.039, Fig. 2d, dataset B: *F* _(3, 112)_ = 0.197, *p* = 0.898, η*p^2^* = 0.008, Fig. 2e]. The results indicated that PSS remained relatively stable over time in the two datasets.

The paired sample t-test showed that the trial numbers between the two responses were insignificant in the two datasets [dataset A: auditory first trial numbers: 345 ± 82 (mean ± SD), visual first trial numbers: 385 ± 90, *t* _(38)_ = −1.531, *p* = 0.134, Cohen’s d = −0.494; dataset B: auditory first trial numbers: 156 ± 50 (mean ± SD), visual first trial numbers: 141 ± 45, *t* _(28)_ = 0.969, *p* = 0.341, Cohen’s d = 0.180].

### 3.2 EEG results

#### 3.2.1 The correlation between power and the individual’s PSS

Prestimulus cortical excitability predicted the individual’s PSS in previous studies. We used two datasets to validate the results. We computed Pearson’s correlation between the prestimulus power difference (Afirst-Vfirst) and the individual’s PSS. This test revealed two significant negative clusters in a time window ranging from −130 to −100 ms around the beta band (25-30 Hz) over left and right temporal sensors (the first cluster value = −415.075, *p* = 0.026; the second cluster value = −304.870, *p* = 0.046; Fig. 3a) in dataset A. We extracted and computed the average power difference within the significant clusters and performed Pearson’s correlation (left-tailed) between the prestimulus power difference (Afirst-Vfirst) and the individual’s PSS in each subgroup. The results indicated that the correlation coefficients were significant except in the group6 (Fig. 3b).

**Fig. 3.**
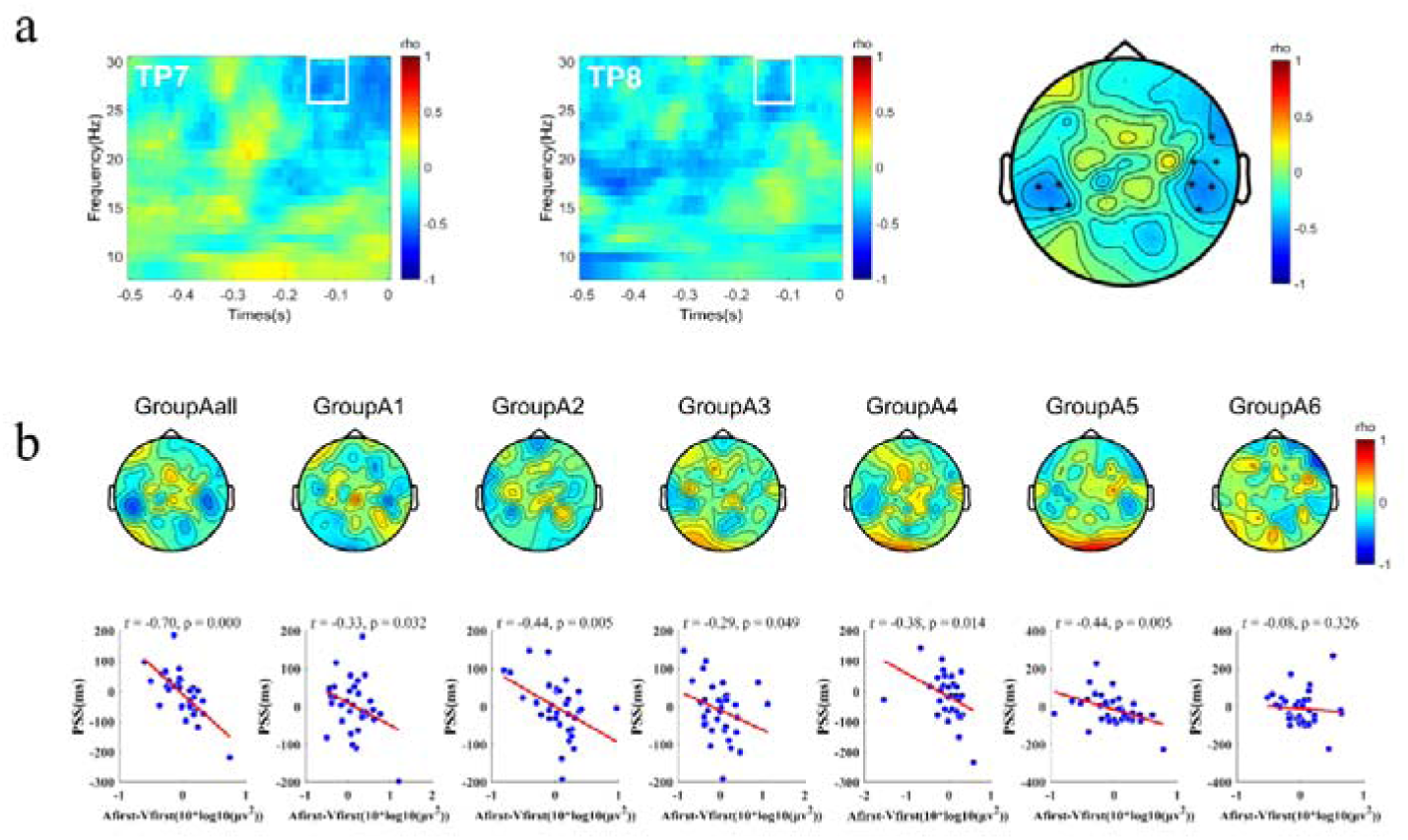
Results of Pearson’s correlation analysis in the beep-flash paradigm. (a) The time-frequency representations of Pearson’s correlations between power difference (Afirst-Vfirst) and the individual’s PSS in the beep-flash paradigm in TP7 channels of the first negative cluster and TP8 channels of the second negative cluster. The white contour line indicated the significant time-frequency cluster. The topography in the right panel shows the average rho map within the time range of −130 to −100 ms and the frequency range of 25 to 30 Hz. Significant channels are highlighted with asterisks. (b) The topography shows Pearson’s correlations between the power difference (Afirst-Vfirst) extracted and averaged within the significant clusters and the individual’s PSS in every subgroup. Below are Pearson’s correlations between the average power difference (Afirst-Vfirst) over significant channels in previous data-driven analysis (i.e., in Fig. 3a) and the individual’s PSS.

The same statistical procedure was used in dataset B. Nonparametric permutation tests revealed a significant negative cluster in a time window ranging from −150 to 0 ms around the beta band (20-25 Hz) over parietal-occipital and occipital sensors (cluster value = −2500.700, *p* = 0.006; Fig. 4a) in dataset B. We extracted and computed the average power difference within the significant clusters and performed Pearson’s correlation (left-tailed) between the prestimulus power difference (Afirst-Vfirst) and the individual’s PSS in each subgroup. The results indicated that the rho is significant only in the groupBall (Fig. 4b).

**Fig. 4.**
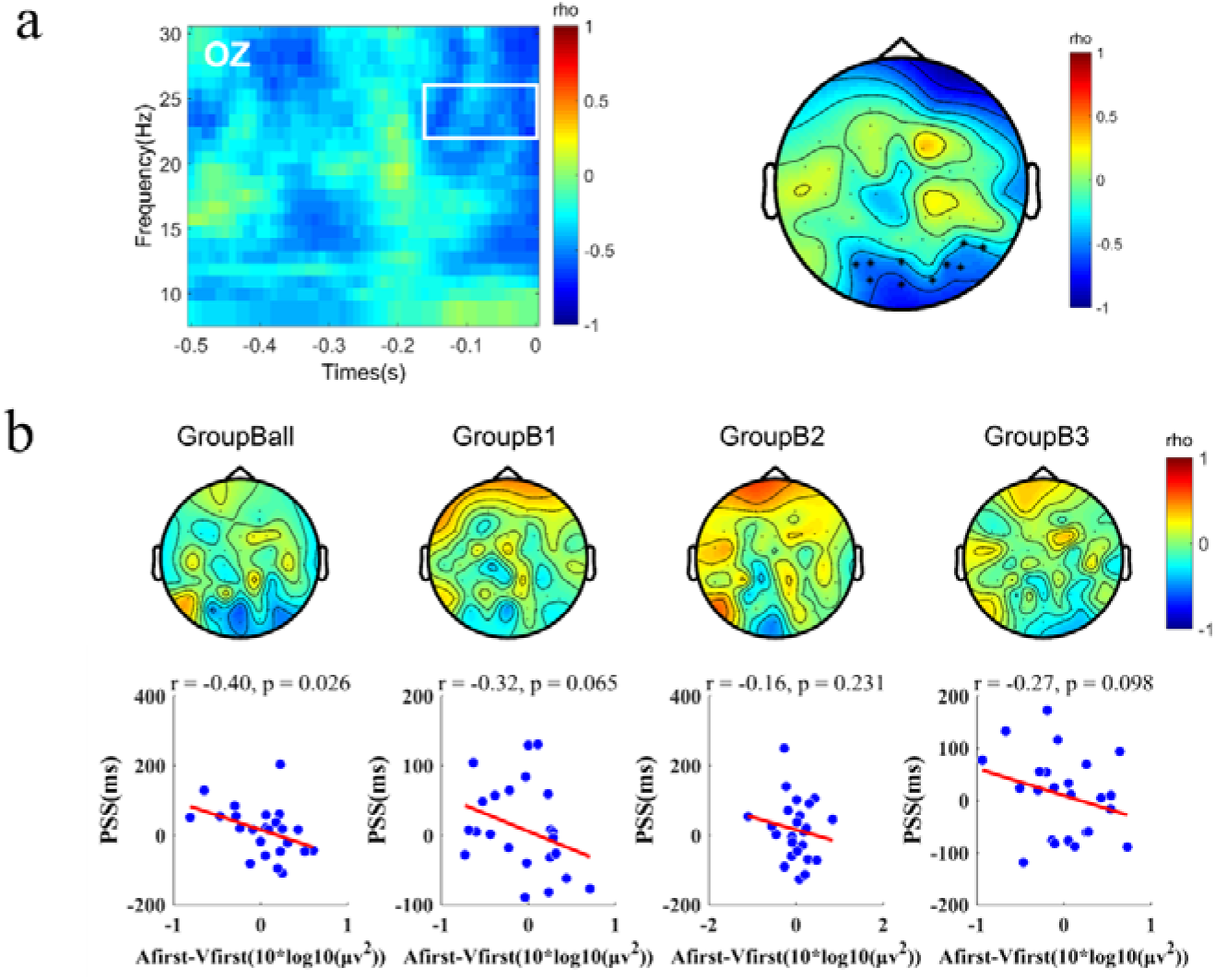
Results of Pearson’s correlation analysis in the stream-bounce paradigm. (a) The time-frequency representations of Pearson’s correlations between power difference (Afirst-Vfirst) and the individual’s PSS in the beep-flash paradigm in OZ channels of the significantly negative cluster. The topography in the right panel shows the average rho map within the time range of - 0.15 to 0 s and the frequency range of 20 to 25 Hz. The white contour line indicated the significant time-frequency cluster. Significant channels are highlighted with asterisks. (b) The topography shows Pearson’s correlations between the power difference (Afirst-Vfirst) extracted and averaged within the significant clusters and the individual’s PSS in every subgroup. Below are Pearson’s correlations between the average power difference (Afirst-Vfirst) over significant channels in previous data-driven analysis (i.e., in Fig. 4a) and the individual’s PSS.

#### 3.2.2 The correlation between dwPLI, power, and the individual’s PSS

Given that the sensory cortical excitability predicted the individual’s PSS ^[11]^ and was modulated by the frontal cortex ^[15]^, we want to test the correlation between the frontal cortex and sensory cortices interactions measured by dwPLI, sensory cortical excitability quantified by power, and the individual’s PSS. First, we explored the correlation between dwPLI difference (Afirst-Vfirst) and power difference (Afirst-Vfirst). We selected Fz, also used in related papers ^[15]^, as the electrode corresponding to the frontal cortex. We picked the channels within the significant clusters as sensory cortical-related channels. We calculated the dwPLI between frontal and sensory cortical-related channels. The functional connectivity matrix’s mean is the final dwPLI estimation. Then, we explored the correlation between dwPLI difference (Afirst-Vfirst) and the individual’s PSS. In the statistical analysis, our primary focus was the time range of −300 to 0 ms and the frequency range of 6-30 Hz. Finally, we investigated the link between dwPLI difference (Afirst-Vfirst), power difference (Afirst-Vfirst), and the individual’s PSS by constructed a simple mediation model.

In dataset A, we investigated the correlations in the two significant clusters separately. In the first cluster, nonparametric permutation tests revealed a significant negative cluster in a time window ranging from −200 to 0 ms over the 6-10 Hz alpha band between dwPLI difference (Afirst-Vfirst) and power difference (Afirst-Vfirst) (cluster value = −150.446, *p* = 0.044; Fig. 5a). In addition, a significant positive cluster was found in a time window ranging from −200 to 0 ms over the 6-10 Hz alpha band between dwPLI difference (Afirst-Vfirst) and the individual’s PSS (cluster value = 186.102, *p* = 0.022; Fig. 5a). The results of mediation analysis showed that the correlation between dwPLI difference and the individual’s PSS could be partially mediated by the power difference (indirect effect: a1b1 = 0.221, 95% CI [0.018 0.476]; direct effect: c’= 0.343, 95% CI [0.066 0.620]; Fig. 5a). In the second cluster, nonparametric permutation tests revealed no significant cluster between dwPLI difference and power difference (cluster value = −22.420, *p* = 0.969; Fig. 5b). There is also no significant cluster between dwPLI difference and the individual’s PSS (cluster value = 28.531, *p* = 0.769; Fig. 5b). Given the consistent and opposing correlation coefficients observed by the eye (Fig. 5b) in the time range of −250 to −150 ms and the frequency range of 26-30 Hz in the two analyses above, we extracted the mean of dwPLI within the time-frequency range for the mediation analysis. The results showed that both the direct and indirect effects were not significant (indirect effect: a1b1 = −0.187, 95% CI [-0.361 0.055]; direct effect: c’= −0.241, 95% CI [-0.050 0.017]; Fig. 5b).

**Fig. 5.**
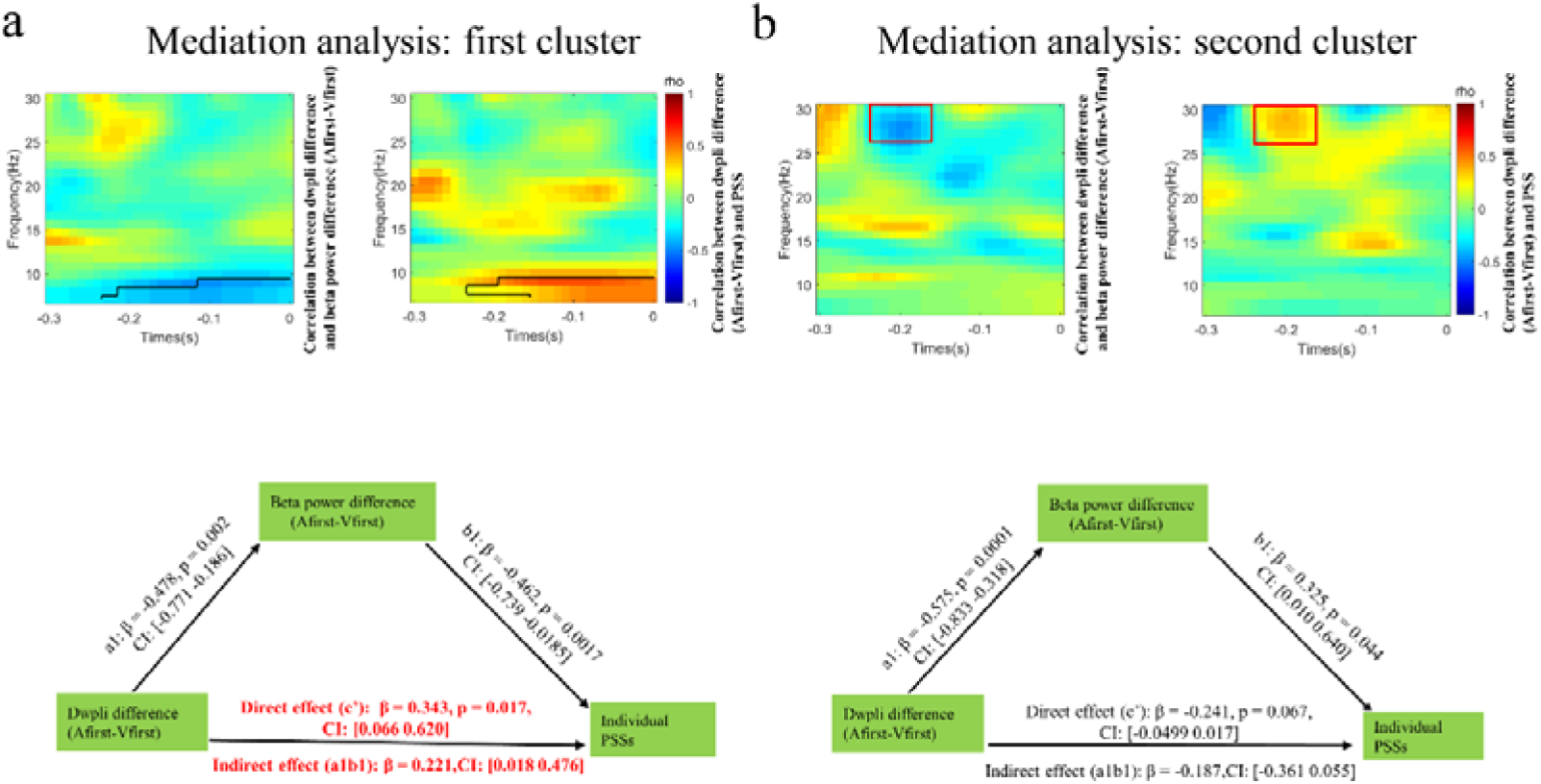
Results of mediation analysis in the beep-flash paradigm. (a) The time-frequency representations of Pearson’s correlations between the dwPLI difference (Afirst-Vfirst) and beta band power difference (Afirst-Vfirst) on the left and the individual’s PSS on the right in the first significant cluster. The black contour line indicated the significant time-frequency cluster. A simple mediation model: spontaneous beta power partially mediated the relationship between prestimulus alpha dwPLI and the individual’s PSS. Red indicates that the path coefficients of direct or indirect effects were significant. (b) The time-frequency representations of Pearson’s correlations between the dwPLI difference (Afirst-Vfirst) and beta band power difference (Afirst-Vfirst) on the left and the individual’s PSS on the right in the second significant cluster. A mediation model: spontaneous beta power did not mediate the relationship between prestimulus alpha dwPLI and the individual’s PSS. Red indicates that the path coefficient is significant. The red contour line indicated the time-frequency cluster selected for mediation analysis.

In dataset B, we investigated the correlations in the significant cluster. Nonparametric permutation tests revealed a significant negative cluster in a time window ranging from −100 to 0 ms over the 14-20 Hz beta band between dwPLI difference (Afirst-Vfirst) and power difference (Afirst-Vfirst) (cluster value = - 141.767, *p* = 0.036; Fig. 6). In addition, a significant positive cluster was found in a time window ranging from −100 to 0 ms over the 14-20 Hz beta band between dwPLI difference (Afirst-Vfirst) and the individual’s PSS (cluster value = 194.509, *p* = 0.010; Fig. 6). The results of mediation analysis showed that the correlation between dwPLI difference and the individual’s PSS could be fully mediated by the power difference (indirect effect: a1b1 = 0.263, 95% CI [0.072 0.532]; direct effect: c’= 0.323, 95% CI [-0.065 0.711]; Fig. 6).

**Fig. 6.**
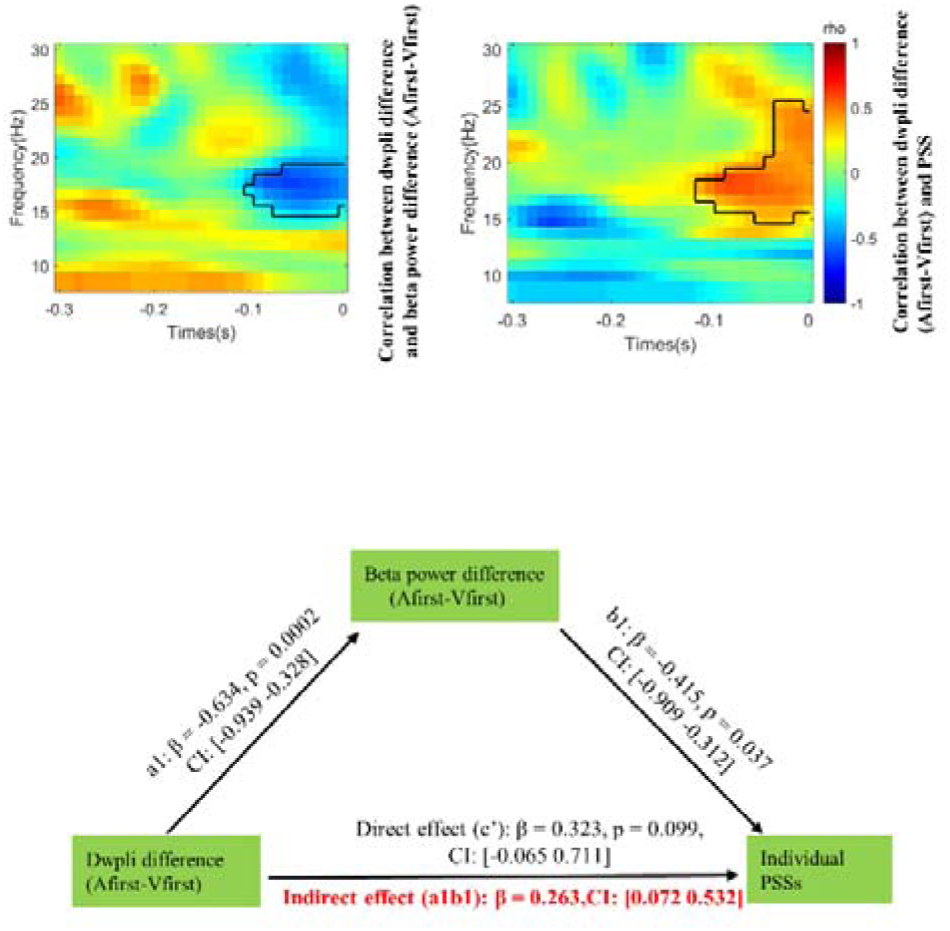
Results of mediation analysis in the stream-bounce paradigm. The time-frequency representations of Pearson’s correlations between the dwPLI difference (Afirst-Vfirst) and beta band power difference (Afirst-Vfirst) on the left and the individual’s PSS on the right. The black contour line indicated the significant time-frequency cluster. A simple mediation model: spontaneous beta power partially mediated the relationship between prestimulus alpha dwPLI and the individual’s PSS. (b) The time-frequency representations of Pearson’s correlations between the dwPLI difference (Afirst-Vfirst) and beta band power difference (Afirst-Vfirst) on the left and the individual’s PSS on the right in the second significant cluster. A parallel mediation model: spontaneous beta power served as a complete mediator in the relationship between prestimulus low beta dwPLI and the individual’s PSS. Red indicates that the path coefficients of direct or indirect effects were significant.

## 4 Discussion

There exist a large number of individual variability in temporal order perception. Previous studies used individually specific stimuli with a single difficulty level to investigate the neural mechanisms underlying individual variability in temporal order perception. We used two experimental paradigms with different visual stimuli to verify them. Our results showed that prestimulus beta band power negatively predicted the individual’s PSS in both experimental paradigms. The brain regions involved in the two paradigms are different. The auditory cortex is involved in the beep-flash paradigm, while the visual cortex is involved in the stream-bounce paradigm. Furthermore, the beta band power was modulated by the functional connectivity of frontal and temporal or occipital areas. Finally, the simple mediation models indicated that sensory cortical excitability could partially or fully mediate the relation between the frontal cortex and individual variability in temporal order perception.

### 4.1 Prestimulus beta power can predict the individual’s PSS

This study found that spontaneous beta band power difference (Afirst-Vfirst) negatively predicted the individual’s PSS in both experimental paradigms. Specifically, the beta band (25-30 Hz) power difference of bilateral temporal channels was negatively related to the individual’s PSS in the beep-flash paradigm, while the beta band (20-25 Hz) power difference of parietal-occipital and occipital channels was negatively correlated with the individual’s PSS in the stream-bounce paradigm. Interestingly, the electrodes corresponding to this effect differ in the two paradigms. At the same time, our experimental results are inconsistent with previous studies ^[10, 11]^ regarding frequency bands and the directionality of correlation.

The bilateral temporal channels might be specifically associated with the auditory cortex, whereas the parietal-occipital and occipital channels should correspond to the visual cortex. Based on the previous hypothesis, auditory and visual cortices contributed to the individual’s PSS ^[11]^. When the excitability of the auditory cortex increases while the excitability of the visual cortex decreases, participants are more sensitive to the auditory stimuli and insensitive to the visual stimuli. Participants are more likely to perceive auditory stimuli as occurring first (i.e., a smaller PSS) ^[11]^. However, the author only observed the effects of the auditory cortex and attributed to most participants’ PSS being negative (auditory stimuli occurring first) ^[11]^. The explanation might be invalid in our study because the numbers of participants with positive PSSs (nineteen subjects) and negative PSSs (twenty subjects) were roughly equal in the beep-flash paradigm. We thought that the auditory cortex might dominate the cross-modal temporal perception. First, the auditory cortex influences the time estimation of visual and auditory stimuli ^[27]^. Second, auditory but not visual ERP predicted the magnitude of the audiovisual TBWs ^[6]^. But this explanation is not applicable in the stream-bounce paradigm. We speculated that a possible reason is that participants demanded more visual attention in the stream-bounce paradigm. A stationary flash is usually mislocalized as lagging behind a moving object in spatiotemporal alignment, referred to as the flash-lag effect (FLE) ^[28]^. Moving objects need more visual attention than static objects ^[29]^. We could speculate that compared to flash stimuli, the perception of the timing of moving dots requires more visual attention, which was also supported by the significantly larger PSS in the stream-bounce paradigm in this study. Taken together, we concluded that the auditory and visual cortex modulated the individual’s PSS depending on task demands induced by the experimental stimuli.

In terms of frequency bands, this study found that the power in the beta band, rather than the alpha band power, is correlated with the individuals’ PSSs. The present beta results seem to disagree with other studies in which alpha band power predicted the individual’s PSS ^[10, 11]^. But previous research also found spontaneous beta band effects for similar stimuli to ours ^[30, 31]^. For example, increased prestimulus beta power in the left superior temporal gyrus (STG) was associated with more illusory perceptions in the auditory-induced double flash illusion (DFI) paradigm using beep-flash stimuli ^[30]^. In addition, subjects with higher prestimulus beta band synchronization between frontal, parietal, and occipital areas were more likely to make bounce rather than stream judgments in the stream-bounce paradigm ^[31]^. Therefore, we thought that our findings were reasonable. Spontaneous alpha band power in the auditory or visual cortex was associated with auditory or visual cortical excitability ^[32, 33]^. Two recent memory-related papers ^[34, 35]^ collectively suggested that the beta power in auditory and visual cortices also played a role in indexing sensory cortical excitability. We thought the beta band power in this study might be correlated with the auditory cortical excitability in the beep-flash paradigm and visual cortical excitability in the stream-bounce paradigm.

Unlike previous research ^[10, 11]^, spontaneous power could negatively predict the individual’s PSS in our study. The different levels of difficulty in stimuli might explain this difference. The individual-specific PSS is relatively challenging for each subject in the first study ^[11]^. Prestimulus alpha power in the auditory cortex positively predicted the individual’s PSS, reflecting that the subjects tried to match the individual temporal order bias. In contrast, the individual-specific JND is relatively easy in the second study ^[10]^. Prestimulus alpha power in the frontal cortex might play a role in overcoming individual temporal order bias. The stimuli in this study varied in levels of difficulty. Based on the Temporal Renormalization Theory ^[5, 36]^, the audiovisual PSS is approximately veridical across different measures and stimuli. The participants might overcome individual temporal order bias to ensure a close match of PSS with the physical time interval of 0 ms in each task. Therefore, we thought that the subjects overcame individual temporal order biases by modulating the excitability of the perceptual cortex in our study.

### 4.2 Spontaneous beta power is a relatively stable marker for predicting the individual’s PSS

The results of Pearson’s correlations between subgroups (Fig. 2c,e) indicated that participants’ PSS is relatively stable, consistent with previous findings ^[4]^, suggesting that temporal order perception is an inherent bias in individuals. Importantly, we also found that spontaneous beta band power reliably predicts individual PSS across all subgroups (Fig. 3b; Fig. 4b). It is worth noting that the stability of the relation between the spontaneous beta power and PSS is stronger in the beep-flash paradigm compared to the stream-bounce paradigm, possibly due to the greater difficulty of the stream-bounce paradigm.

### 4.3 Frontal cortex affected individual PSSs by modulated sensory cortical excitability

Many researchers have found that the frontal cortex affects visual cortical excitability, leading to better task performance in visual detection ^[15]^ and discrimination tasks ^[17]^. Some studies also found that the frontal cortex could modulate auditory cortical excitability in humans ^[37]^ and nonhuman primates ^[38]^. A recent review suggested that at least two similar attention networks exist for visual and auditory modalities ^[39]^. We found that the functional connectivities of the frontal-left-temporal cortex in the alpha (6-10 Hz) band could shape the beta band power in the temporal cortex in the beep-flash paradigm. At the same time, the functional connectivity of the frontal-occipital cortex in the beta (14-20 Hz) band could modulate the beta band power in the occipital cortex in the stream-bounce paradigm. Our results indicated that the frontal cortex could modulate the temporal and occipital cortical excitability, consistent with the previous results.

Interestingly, we found that the functional coupling between the frontal and sensory cortices could affect the excitability of sensory cortices and further modulate the individual’s PSS. These results specified the link between frontal top-down modulation, sensory cortical excitability, and behavioral performance in perceptual tasks. Notably, auditory cortical excitability might only partially explain behavioral performance modulated by the frontal cortex in the beep-flash paradigm. Given that the prestimulus local and network activities could jointly and concurrently influence perceptual performance ^[30, 40]^, we thought our results were reasonable. We guessed that frontal-temporal functional coupling might also be associated with auditory attention.

## 5 Conclusion

Our study used two experimental paradigms where the subjects performed the temporal order judgment (TOJ) task to verify the correlation between spontaneous neural oscillations and the individual’s PSS found in previous studies. We found that the ongoing beta band power in the auditory cortex was negatively correlated with the individual’s PSS in the beep-flash paradigm while positively related to the individual’s PSS in the stream-bounce paradigm. Furthermore, the functional connectivity between the frontal and auditory or visual cortex could modulate the spontaneous neural oscillation power and further shape the individual’s PSS. We could draw three conclusions from the above results. First, spontaneous neural oscillation in sensory cortices indexed cortical excitability and could predict individual variations in temporal order perception. Second, the visual or auditory cortex could play a role in reflecting the individual’s temporal order perception depending on different experimental paradigms. Finally, the frontal cortex could modulate sensory cortical excitability and further shape the individual’s temporal order perception.

## Acknowledgments

This work was supported in part by the National Key Research & Development Program of China (No. 2022YFF1202400) and the National Natural Science Foundation of China (No.62276181).

